# Chemical Stimulation Sustains Bioluminescence of Living Light Materials

**DOI:** 10.1101/2025.07.16.664986

**Authors:** Giulia Brachi, Jessica McKean, Cheng Pau Lee, Joy Edwin-Ezeh, Wil V. Srubar

**Affiliations:** Department of Civil, Environmental, and Architectural Engineering, 1111 Engineering Drive, University of Colorado Boulder, Boulder, Colorado USA 80309; Materials Science and Engineering Program, 4001 Discovery Drive, University of Colorado Boulder, Boulder, Colorado USA 80303

## Abstract

Bioluminescence offers a powerful tool for real-time, label-free sensing for living materials. However, conventional approaches often rely on mechanical stimulation, which is difficult to standardize, spatially localize, and sustain over time. Here, we introduce a chemical strategy to stimulate and sustain bioluminescence in the marine dinoflagellate *Pyrocystis lunula*, enabling the fabrication of robust, adaptive, light-emitting living materials. By embedding *P. lunula* into 3D-printed, ionically crosslinked alginate hydrogel scaffolds, we engineered architecturally stable living materials with long-term cellular retention, viability, and light-emitting capacity. Exposure to acidic and basic environments enabled chemically resolved sensing and response *via* distinct bioluminescent signatures: acid triggers intense, localized, and persistent emission up to 25 minutes, while base induces a diffuse, biphasic emission indicative of cellular stress. Notably, coupling chemical with mechanical stimulation yields a synergistic enhancement of bioluminescence, achieving significantly greater amplitude and duration of light emission without compromising cell reactivity. Longitudinal studies over four weeks demonstrated that our living-light materials retain responsiveness and structural integrity across repeated stimulation cycles, overcoming the limitations of single-use mechanical activation. Together, these findings establish a robust new platform for programmable, light-emitting living materials with applications in biosensing, soft robotics, and environmental monitoring.

## Main

Biological light emission offers a powerful, self-sustained strategy for real-time sensing for applications in biosensing, soft robotics, and environmental monitoring.^1–5^ Among living systems, bioluminescent microorganisms capable of converting environmental cues directly into optical signals are particularly well-suited for autonomous biosensing.^6–9^ Dinoflagellates, such as *Pyrocystis lunula*, stand out for their ability to produce intense, rapid flashes of light in response to mechanical stimulation, an evolved defense mechanism against predators in turbulent marine environments.^10–13^ This intrinsic mechanoresponsiveness has enabled the use of dinoflagellates as highly sensitive reporters for transient mechanical inputs and fluidic perturbations.^14–17^ However, mechanical stimulation is inherently transient, difficult to spatially or temporally control, and often causes irreversible structural degradation, limiting reusability in engineered living devices that require sustained light output.^18^ Chemical stimulation offers a promising alternative, enabling tunable and reproducible bioluminescence through defined perturbations of the cellular environment.^19–24^ However, its integration as a reliable activation mechanism within engineered bioluminescent systems remains largely unexplored.

Here, we develop a living materials platform by encapsulating *P. lunula* within 3D-printed, ionically crosslinked alginate scaffolds, enabling responsive bioluminescent matrices capable of sustained light emission in response to chemical stimuli. We show that chemically defined cues, specifically acidic and basic environments, induce divergent and tunable bioluminescent signatures, which are retained in both free-swimming and hydrogel-confined states. High-resolution N-STORM microscopy and IVIS imaging substantiate that acid stimulation generates intense and localized emission, while basic conditions elicit a biphasic, cell-wide response indicative of organelle degradation and cell death. Implementation of a dual-mode activation strategy that couples chemical and mechanical stimulation results in synergistically enhanced light output, revealing intracellular potentiation mechanisms that amplify native mechanosensitivity.^25^ Furthermore, we show that chemical stimulation enables repeated, long-term activation cycles without compromising viability, outperforming conventional mechanically triggered systems that typically suffer from irreversible structural degradation and consequent functional loss after single use.^26^ Together, these findings establish a robust and reconfigurable platform for sustained light-emitting materials, with potential applications in biosensing, soft robotics, and environmental monitoring.^5,27,28^

### Modulation of Bioluminescence in *Pyrocystis lunula* through Chemical Stimulation

To establish chemical stimulation as a viable mechanism for sustained bioluminescent actuation, *P. lunula* cells were exposed to controlled acidic (pH 4) and basic (pH 10) environments, and their responses evaluated using complementary qualitative and quantitative imaging techniques (Fig. 1a).

**Figure 1.**
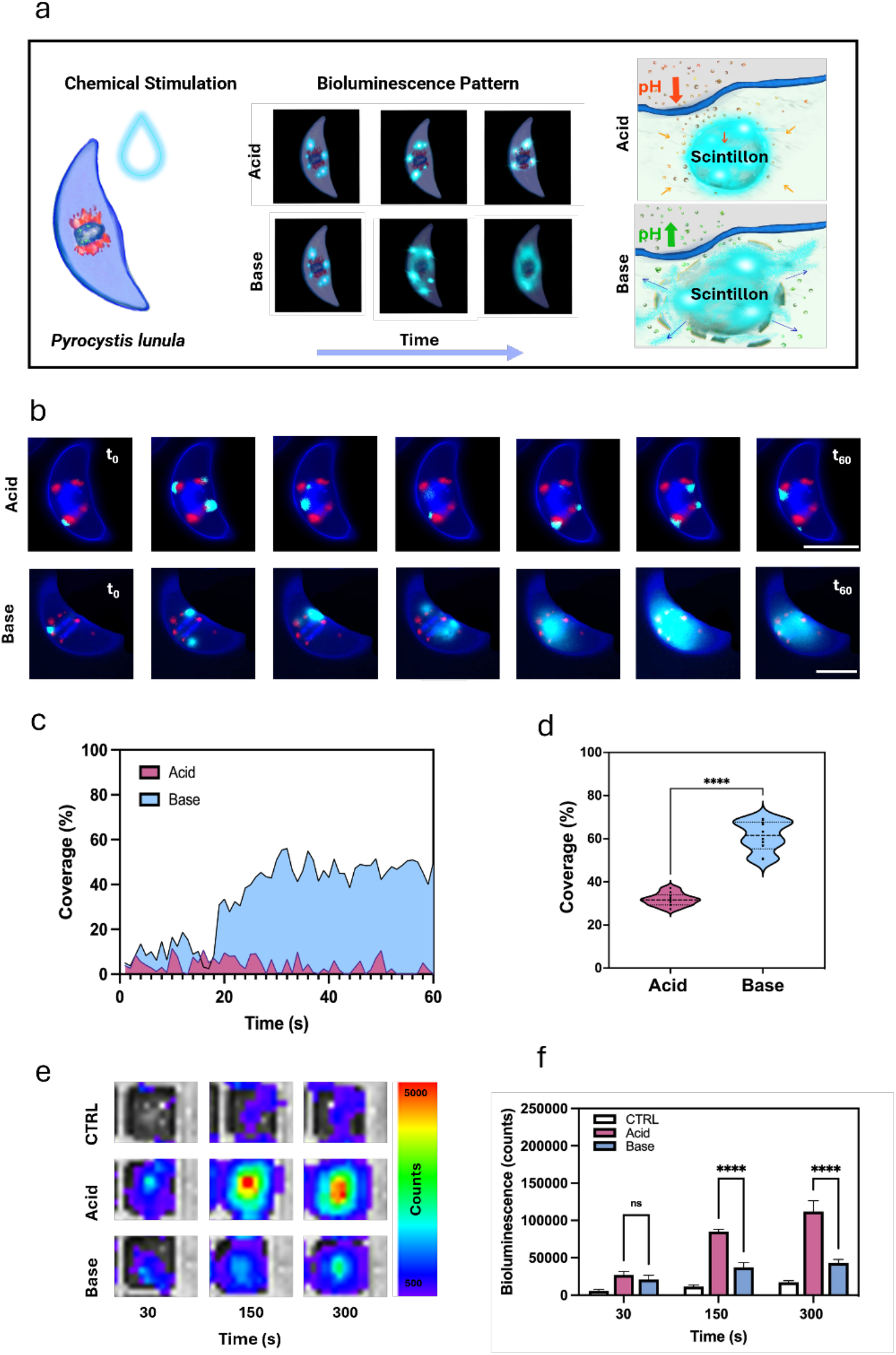
Chemical stimulation of bioluminescence in *Pyrocystis lunula*. a, Schematic of pH-dependent light emission. Cells exposed to acidic (pH 4) or basic (pH 10) environments were analyzed by fluorescence microscopy and IVIS imaging. Acidic treatment triggered localized, intracellular bioluminescence, while basic conditions induced diffuse, widespread emission. Cross-sectional views highlight localized scintillons in acidic conditions and broader emission patterns in basic conditions. b, N-STORM images showing distinct emission morphologies in acid-treated (top) and base-treated (bottom) cells. Scale bars, 50 μm. c, Quantification of bioluminescent area reveals larger emission domains under basic conditions; acid-treated cells shown in pink, base in blue. d, Violin plots summarizing bioluminescence coverage across individual cells (N = 10), indicating statistically distinct responses. e, IVIS photon flux heatmaps at 30, 150, and 300 s illustrate greater spatial uniformity in acidic conditions and increased heterogeneity in basic conditions. f, Total photon flux quantification (N = 6) confirms significantly enhanced bioluminescence under acidic stimulation at later timepoints. Data are mean ± s.d.; two-way ANOVA with Tukey’s post hoc test, ****P < 0.0001; ns, not significant.

Super-resolution N-STORM microscopy revealed distinct bioluminescent signatures at the single-cell level under acidic and basic conditions, whereas no signal emission was detected in control cells in standard L1 medium (Fig. 1b; Fig. S1a-b). Acid-treated cells exhibited intense, spatially confined emission localized to intracellular compartments consistent with scintillons. This is in agreement with early macroscale studies from the 1970s showing that direct external acidification mimics mechanical stimulation by driving proton flux and scintillon acidification (Fig. 1b, top row).^12,29^ In contrast, base-stimulated cells exhibited a biphasic emission profile: an initial focal signal originating from scintillons was followed by a diffuse, cell-wide luminescent response (Fig. 1b, bottom row). This transition is attributed to progressive intracellular ionic imbalance compromising the membrane integrity with consequent release of luminescent compounds throughout the cell.^30^

Interestingly, the two stimulation modes elicited markedly different spatial and temporal bioluminescence profiles of light emission (Fig. 1c). Acid treatment produced sustained, spatially confined emission localized to discrete intracellular regions, covering 3.8 ± 3.4% of the total cell area at each measured timepoint. In contrast, base stimulation started with a similarly localized signal (i.e., 4.9%) but rapidly transitioned to a diffuse, cell-wide luminescent response reaching 50.0% by 60 s. When emission area was quantified across the full temporal sequence (N=10), acid-treated cells exhibited a cumulative coverage of 31.8 ± 2.9% of total cell area, while base-treated cells showed significantly higher, diffuse whole-cell coverage of 61.2 ± 7.0% (Fig. 1d).

To capture bioluminescence dynamics across large cell populations, quantitative IVIS imaging was performed over a 300-second period in wells containing ∼2.5 × 10^3^ freely suspended *P. lunula* cells (N = 6). Heatmaps revealed that acid exposure elicited a markedly higher and more temporally sustained bioluminescent output relative to base-treated samples (Fig. 1e), with signal stability maintained over several minutes. Quantitative analysis confirmed higher peak intensity (acid: 112000 ± 14315 vs. base: 43000 ± 4729 photon counts) for the acid group (Fig. 1f), consistent with the greater physiological tolerance of *P. lunula* to mildly acidic conditions relative to alkaline stress. ^31^

Together, these results demonstrate that chemically defined cues can be used not only to initiate bioluminescence, but also to modulate its spatial and temporal profile, thus offering a tunable and programmable strategy for engineering responsive light behavior in living systems.

### Encapsulation of *P. lunula* in Biocompatible Hydrogel Matrices

To integrate chemically responsive dinoflagellates into functional living materials, we developed an encapsulation strategy that maintains cellular viability, enables molecular exchange, and ensures spatial confinement. *P. lunula* cells were embedded in 4 wt% alginate and ionically crosslinked to form stable hydrogel constructs (Fig. 2a).^32,33^

**Figure 2.**
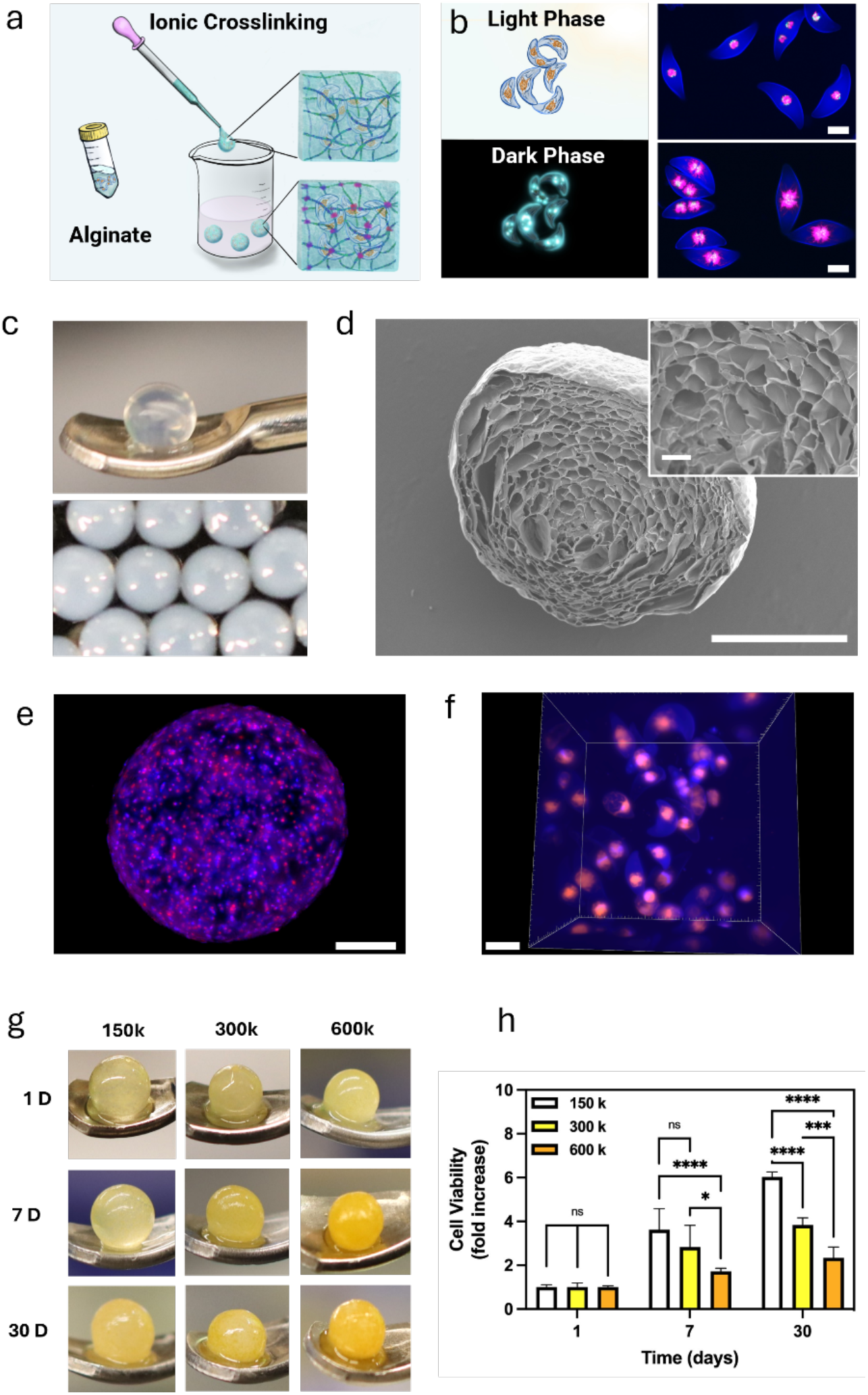
Encapsulation of *Pyrocystis lunula* in alginate hydrogels enables long-term viability and uniform spatial confinement. a, Schematic of the encapsulation process, in which *P. lunula* cells are suspended in sodium alginate and ionically crosslinked in CaCl_2_ to form stable hydrogel bead constructs. b) Fluorescence micrographs of circadian-synchronized cultures under light and dark phases. Chloroplast (autofluorescence, Cy5, magenta), cytoplasm (DAPI, blue); Scale bar, 50 μm. c, Optical images of individual and grouped alginate beads (diameter = 1.6 ± 0.1 mm). d, Scanning electron micrograph (SEM) of a lyophilized bead cross-section. Main scale bar, 1 mm; inset, 100 μm. e, Fluorescence imaging of DAPI-stained cryosections showing homogeneous cell distribution. Scale bar, 500 μm. f, Volumetric 3D light-sheet microscopy further confirms uniform dispersion with no sedimentation. Scale bar, 80 μm. g, Brightfield images of beads cultured for 30 days reveal chlorophyll changes indicative of cell proliferation. Scale bar, 80 μm. h, Quantification of cell viability over time using MTS assay for three initial seeding densities (150k, 300k, and 600k cells mL^−1^). Data are mean ± s.d. (n = 4). Statistical significance determined by two-way ANOVA with Tukey’s post hoc test (*P* < 0.05 to \*\*\**P* < 0.0001).

Prior to encapsulation, *P. lunula* cultures were conditioned under inverted light/dark conditions (light: 5 pm-7 am; dark: 7 am-5 pm), as confirmed by chloroplast migration patterns indicative of circadian reprogramming (Fig. 2b), consistent with recent observations by Schramma *et al*.^34^ Hydrogel beads (1.6 ± 0.1 mm diameter) were formed by dispensing sodium alginate into a calcium chloride solution (Fig. 2c, S2a,b). Scanning electron microscopy (SEM) of bead cross-sections revealed an interconnected porous network (Fig. 2d) that facilitates nutrient and gas exchange while physically retaining cells within the matrix, providing a structurally supportive environment for sustained biological activity.^35^

Uniform spatial distribution of cells within the hydrogel matrix is critical for consistent bioluminescent output across living devices.^36^ To achieve this, *P. lunula* cells were gently homogenized into the alginate precursor prior to crosslinking. Fluorescence imaging of cryosectioned beads (Fig. 2e) and volumetric light-sheet microscopy (Fig. 2f) confirmed even cellular dispersion. No aggregation or sedimentation was observed, ensuring functional uniformity across devices.

To assess biocompatibility and proliferative potential within the encapsulated environment, hydrogel beads were fabricated at three distinct initial seeding densities (150, 300, and 600 × 10^3^ cells mL^−1^). Brightfield imaging revealed dynamic yellow color shifts in the chloroplasts over time (Fig. 2g), reflecting cellular proliferation. Quantitative viability analysis using MTS metabolic assays over a 30-day culture period (Fig. 2h) demonstrated long-term viability across all tested conditions. Interestingly, beads seeded at 150 × 10^3^ cells mL^−1^ showed the greatest MTS signal increase (6-fold by day 30, *P* < 0.0001), suggesting enhanced proliferative capacity under less crowded conditions, while higher initial densities reached a plateau, likely due to space limitations imposed by the hydrogel microenvironment.^37,38^

Collectively, these results establish alginate as a robust matrix for *P. lunula* encapsulation, supporting metabolic activity and proliferation while preserving spatial uniformity, which are key parameters for engineering scalable, light-emitting living materials.

### Bioprinting of Chemically Responsive Living Light Materials

To enable scalable fabrication of bioluminescent living materials, we optimized dinoflagellate-laden alginate formulations for high-throughput 3D bioprinting (Fig. 3a).

**Figure 3.**
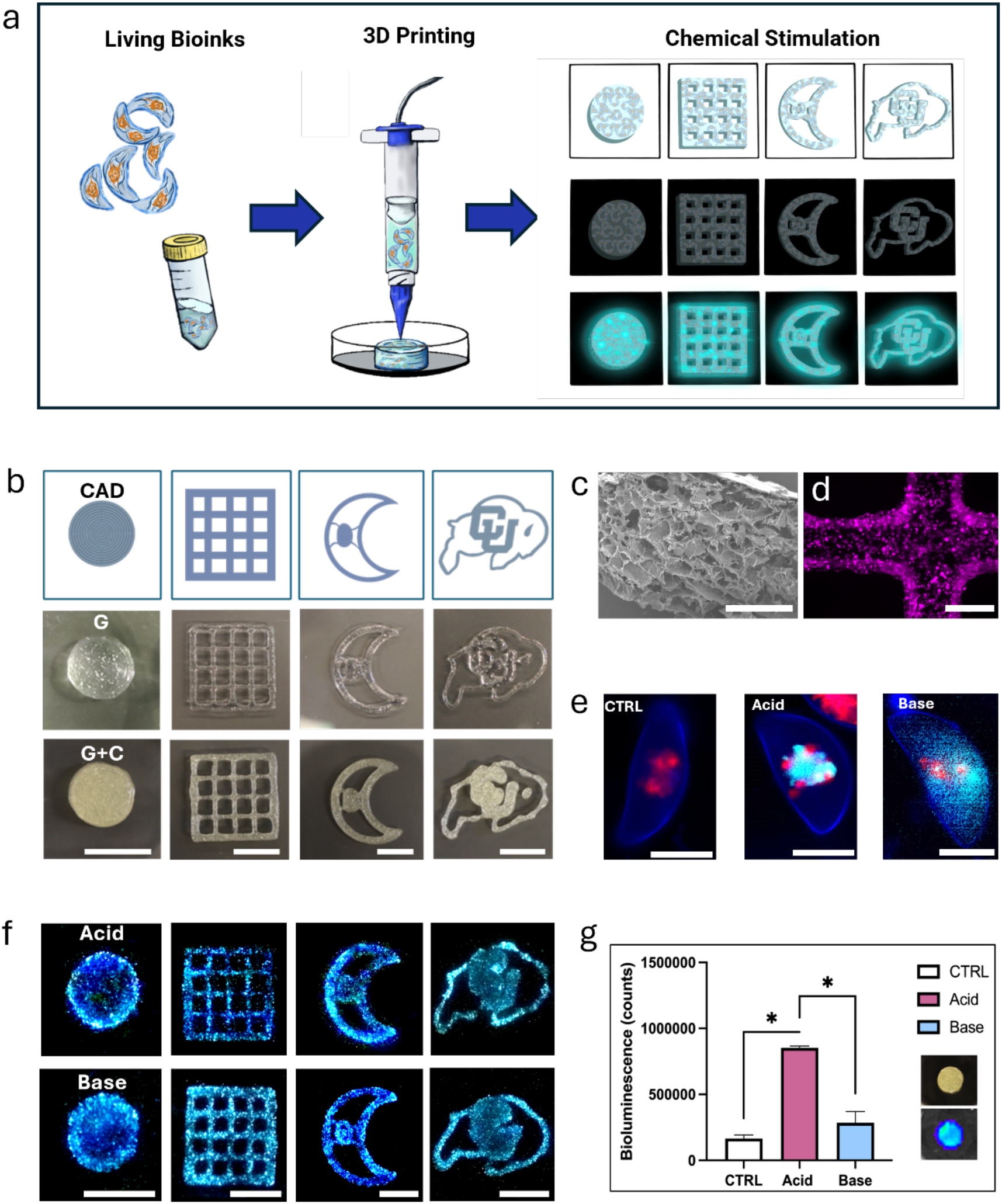
3D bioprinting of *Pyrocystis lunula*–laden alginate bioinks enables spatially defined, chemically stimulated bioluminescence. a, Schematic of the bioprinting workflow. *P. lunula* cells were suspended in sodium alginate, partially pre-crosslinked with CaCl_2_, and printed into defined geometries prior to chemical stimulation. b, CAD renderings (top) and brightfield images of 3D-printed constructs without (G = Gel) or with embedded cells (G+C = Gel + Cells). Scale bars, 10 mm. c, Scanning electron microscopy (SEM) of cryosectioned constructs showing an interconnected porous microarchitecture. Scale bar, 1 mm. d, Fluorescence imaging confirms uniform distribution of *P. lunula* (magenta, Cy5) throughout the 3D-printed constructs. Scale bar, 750 μm. e, N-STORM imaging of encapsulated *P. lunula* in control, acidic, and basic conditions. Scale bars, 50 μm. f, Brightfield images of 3D-printed constructs following acid (top) or base (bottom) stimulation. Scale bars, 10 mm. g, Quantification of IVIS luminescence reveals significantly greater signal under acid stimulation (mean ± s.d., n = 8; one-way ANOVA, *P* < 0.05).

Printability was optimized by tuning the bioink viscosity through partial ionic pre-crosslinking. Alginate solutions (4 or 6 wt%) were combined with CaCl_2_ at defined volumetric ratios, ranging from 1:0 (no pre-crosslinking) to 4:1 and 1:1 (alginate to CaCl_2_), using a dual-syringe mixing system (Fig. S3a). Bioinks were then extruded and evaluated for shape fidelity (Fig. S2b), with optimal performance achieved using 4 wt% alginate at a 4:1 alginate:CaCl_2_ ratio, yielding stable, high-resolution grids in both control and cell-laden prints (Fig. 3b). Rheological analysis of the optimized alginate formulation confirmed its suitability for extrusion-based bioprinting. Strain sweep measurements showed a dominant G′ (845 ± 16 Pa) over G″ of (96 ± 2 Pa) at 0.1–1% strain, indicating stable viscoelastic behavior for shape retention during and after printing (Fig. S3c).^39,40^ Flow sweep analysis showed shear-thinning behavior, with viscosity decreasing steadily from 1.36 to 0.21 Pa·s across 0.1–100 s^−1^ (Fig. S2d).^41^ This enables smooth extrusion and rapid recovery, essential for preserving print fidelity in high-resolution bioprinting.^42,43^

As with bead-based encapsulation, SEM imaging of cryosectioned constructs revealed a porous architecture supporting nutrient exchange and cellular confinement (Fig. 3c). Fluorescence microscopy confirmed that the printing process preserved *P. lunula* viability, with uniform cell distribution and strong Cy5 signal throughout the hydrogel (Fig. 3d), along with evidence of *in situ* proliferation (Fig. S3e).

To assess biosensing function, constructs were then exposed to acidic or basic environments and evaluated at both the micro-and macroscale (Fig. 3e, f). N-STORM imaging of encapsulated dinoflagellates exposed to acidic or basic environments revealed bioluminescence profiles consistent with those in suspension cultures, indicating that hydrogel confinement does not impair stimulus responsiveness (Fig. 3c). Macroscopic IVIS imaging showed a 2.4-fold increase in luminescence under acidic conditions compared to base-treated scaffolds (P < 0.05), consistent with the known acid tolerance of *P. lunula* (Fig. 3f), and extended signal emission up to 25 minutes after stimulation (Fig. S3f).^6^

These findings demonstrate that the bioprinted living light materials retain full biosensing functionality and respond predictably to chemical stimulation.

### Synergistic Enhancement of Bioluminescence via Chemo-Mechanical Activation

To amplify the light-emitting capacity of dinoflagellate-laden constructs, we implemented a dual-mode activation strategy combining chemical with mechanical stimulation (Fig. 4a). This builds on the established mechanosensitive bioluminescent response of *P. lunula*, which emits light rapidly in response to shear or compressive stress.^5,17,44^

**Figure 4.**
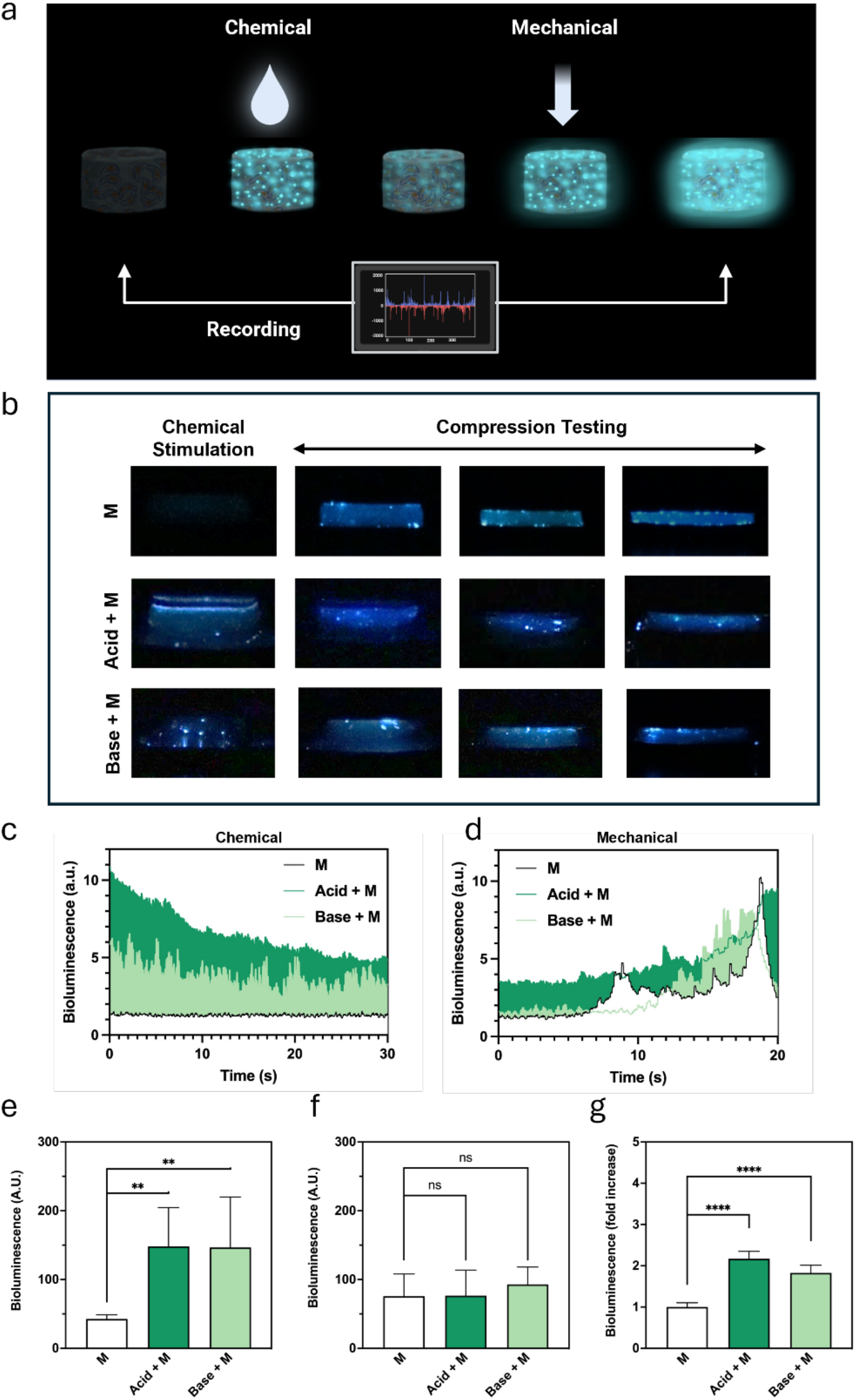
Synergistic enhancement of bioluminescence *via* combined chemical and mechanical stimulation. a, Schematic illustrating dual-mode activation of dinoflagellate-laden 3D-printed hydrogel constructs through chemical stimulation (acid or base) followed by mechanical compression. *Created in BioRender.* b, Time-lapse bioluminescence images of 3D-printed constructs under mechanical-only (top), combined acid + mechanical (middle), and base + mechanical (bottom) stimulation. c, Representative light emission profiles over 30 s following chemical stimulation alone. d, Representative light emission profiles during mechanical compression, showing preserved mechanoresponsiveness across all groups. e, Quantification of cumulative emitted bioluminescence (a.u.) after chemical stimulation. f, Quantification of cumulative emitted bioluminescence (a.u.) during mechanical compression after chemical stimulation. g, Quantification of overall emitted bioluminescence (fold increase) showing that combined chemical and mechanical stimulation yields a synergistic increase in total light output. Data are mean ± s.d. (n = 8); significance determined by two-way ANOVA with Tukey’s post hoc test, P < 0.05.

3D-printed hydrogel scaffolds were chemically preconditioned at pH 4 or pH 10 and subsequently subjected to compressive loading during real-time bioluminescence imaging. As shown in Fig. 4c, both acid and base treatments significantly enhanced spontaneous light emission relative to untreated controls, with acid-primed samples exhibiting the most pronounced peak intensity. Upon mechanical compression (Fig. 4d), all groups showed comparable emission profiles, indicating that chemical conditioning preserved the native mechanosensitive response.

Quantification of the cumulative area under the emission curves revealed that chemical stimulation alone led to an approximately 3.5-fold increase in total luminescence compared to controls (acid: 148.3 ± 56.3 a.u.; base: 146.6 ± 73.4 a.u.; Fig. 4e). In contrast, mechanical activation alone produced similar light outputs across all groups (Fig. 4f), further confirming that preconditioning does not impair mechanoresponsiveness.

Notably, the combination of chemical stimulation and mechanical input resulted in a synergistic enhancement of bioluminescence, with total luminescent output reaching 661.3 ± 118.4 a.u. for acid-treated constructs and 556.3 ± 104.0 a.u. for base-treated ones, compared to 304.6 ± 31.6 a.u. in controls (Fig. S4a). These results are consistent with intracellular reprogramming under pH stress that potentiates mechanosensitive bioluminescence mechanisms, likely by enhancing the sensitivity of scintillon-associated proton channels and cytosolic Ca^2+^ signaling pathways.^13,45^ This potentiation likely accounts for the more-than-two-fold increase in total luminescent output observed upon combined stimulation (Fig. 4g).

Together, these results demonstrate that the light-emitting function of living materials can be modulated through combined chemical and mechanical inputs, offering a versatile route to significantly enhance and sustain light emission in living material systems.

### Cyclic Chemical Stimulation of Living Light Materials

To assess long-term viability, durability, and reusability, *P. lunula*-laden hydrogel constructs were subjected to weekly stimulation under acidic or basic conditions over four weeks (Fig. 5a).

**Figure 5.**
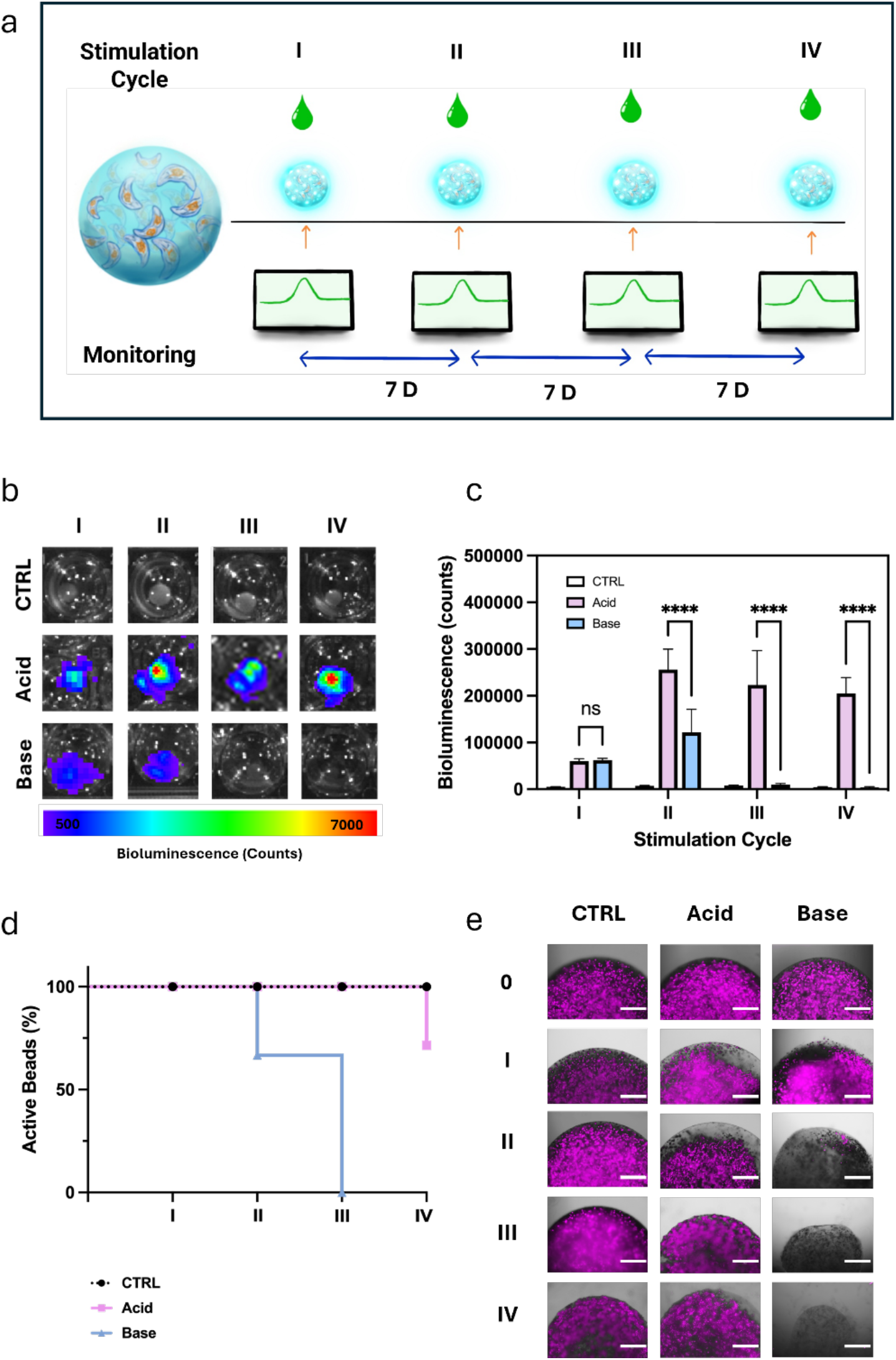
Cyclic chemical stimulation and longevity of living light materials. a, Schematic of a longitudinal stimulation protocol involving weekly exposure to acidic or basic environments and IVIS-based bioluminescence monitoring across four cycles. b, Representative IVIS images illustrating bioluminescence profiles following repeated chemical stimulation. c, Peak photon flux values recorded over time to assess signal retention across treatment conditions (N = 8). d, Kaplan-Meier-like analysis tracking the persistence of functional bioluminescence in 3D-printed constructs over the multi-week stimulation protocol (N = 8). e, Fluorescence imaging (Cy5, magenta) of cell-laden constructs post-stimulation reveals treatment-dependent differences in viability and structural integrity. Scale bars, 750 μm. Data are shown as mean ± s.d.; significance assessed via two-way ANOVA with Tukey’s post hoc test.

IVIS imaging revealed that acid-conditioned constructs maintained robust bioluminescent output across all four cycles, while base-treated samples showed progressive signal attenuation, with complete functional loss by week three (Fig. 5b, c). Quantification of peak bioluminescence revealed sustained enhancement in acid-treated constructs over time (N=8). By the second cycle, peak luminescence reached 256000 ± 43770 a.u. for acid-treated constructs and 122000 ± 49181 a.u. for base-treated ones. Base signal dropped by approximately 92% in the third cycle (9850 ± 2574 a.u.) and another ∼58% by the fourth (4160 ± 1498 a.u.), resulting in a 97% overall decrease. At this point, base-treated samples were indistinguishable from unstimulated controls (4480 ± 1020 a.u.). In contrast, acid-treated constructs maintained high output through all cycles (223000 ± 73868 a.u. at cycle 3; 205000 ± 33803 a.u. at cycle 4), confirming superior and longer-lasting luminescent function (P < 0.0001, two-way ANOVA with Tukey’s post hoc test).^45– 47^

Functional viability was further supported by a Kaplan-Meier-like curve, showing 75% luminescent activity in acid-treated constructs through week 4, compared to 0% reactivity by week 3 in the base-treated group (Fig. 5d). Fluorescence imaging corroborated these findings at the cellular level: acid-treated and control scaffolds maintained dense, viable microalgal populations (evidenced by strong Cy5 signal) and intact hydrogel architecture, whereas base-treated samples exhibited extensive cell loss and structural degradation (Fig. 5e), in line with prior reports of membrane destabilization under alkaline stress.^48^

These results collectively demonstrate that acid stimulation preserves the long-term viability and reusability of living light-emitting materials, enabling repeated and reliable activation for dynamic biosensing applications.

## Discussion

This work establishes chemical stimulation as a robust and repeatable strategy for triggering bioluminescence in *Pyrocystis lunula*, enabling the development of programmable, light-emitting living materials. Embedding dinoflagellate cells within ionically crosslinked, 3D-printed alginate scaffolds enabled the fabrication of robust living devices with sustained cellular retention, proliferation, and structural integrity over time.^49–51^

Living-light materials exhibited predictable responsiveness to defined chemical stimuli, with acidic and basic environments enabling temporally sustained and tunable bioluminescent output at both the micro- and macroscale.^52^ Introducing a dual mode activation strategy combining chemical priming with mechanical compression resulted in a synergistic enhancement of signal output, with chemical preconditioning preserving the platform’s intrinsic mechanoresponsiveness.^53^

Unlike mechanically activated biosensors that undergo structural degradation and functional exhaustion after single-use, chemical stimulation supports repeated activation cycles, demonstrating the robustness and reusability of the constructs for applications requiring sustained or repeated light emission.^54^ Importantly, our results collectively indicate that acidic stimulation, compared to basic conditions, offers superior performance for engineering living light materials due to its higher bioluminescence intensity, cyclic recoverability, and long-term cell viability.

Together, these findings advance a versatile platform for light-emitting living devices with potential applications in adaptive biosensing, soft robotics, and environmental monitoring.^36^ Future work will focus on expanding the stimuli library and integrating multiplexed inputs to enhance control and scalability.^55,56^

## Methods Materials

*Pyrocystis lunula* (strain CCMP731) was obtained from the Bigelow National Center for Marine Algae and Microbiota (East Boothbay, ME, USA). Sea salt (ASTM D1141-98) was purchased from Lake Products Company LLC (Florissant, MO, USA). Sodium nitrate (NaNO3), sodium dihydrogen phosphate (NaH_2_PO4), nickel sulfate (NiSO4), and potassium chromate (K_2_CrO4) were obtained from Thermo Fisher Scientific (Waltham, MA, USA). Sodium alginate, glutaraldehyde solution (25%), 4′,6-diamidino-2-phenylindole dihydrochloride (DAPI) were obtained from Sigma-Aldrich (Burlington, MA, USA). Calcium chloride (CaCl_2_) was purchased from Carolina Biological Supply (Burlington, NC, USA). Sterile 0.22 μm membrane filters (1 L capacity) were from Corning (Tewksbury, MA, USA). CellTiter 96® Aqueous One Solution Cell Proliferation Assay (MTS) was purchased from Promega (Madison, WI, USA). Cell culture wells (6-well and 96-well plates), flasks (250 mL) and ibidi μ-Dish glass-bottom imaging dishes, were from Thermo Fisher Scientific. Standardized pH buffer solutions (pH 4 and pH 10) were obtained from Biopharm Laboratories (Bluffdale, UT, USA). All solvents were of analytical grade.

### Culture of *Pyrocystis lunula*

*Pyrocystis lunula* was cultured in L1 medium composed of 41.95 g L^−1^ ocean salts, supplemented with trace metals and vitamins according to Guillard’s formulation.^28^ Media were sterilized *via* 0.22 µm filtration. Cultures were maintained in sterile 250 mL cell culture flasks under a 12 h light/12 h dark inverted circadian cycle (lights on from 5 pm to 7 am, cool white LED illumination) at 24°C. To ensure physiological stability and exponential growth, cultures were passaged every three weeks at a 1:3 dilution ratio.

### Real-Time Bioluminescence Imaging of Chemically Stimulated Cells

To resolve bioluminescent dynamics at the single-cell level, *P. lunula* cells harvested during their dark phase were centrifuged at 1500 rpm for 10 min and resuspended in sterile L1 medium. After gravitational settling for 5 min in the dark, 20 µL of cell suspension was gently pipetted onto glass-bottom dishes, adding 50 µL of 1:1000 DAPI solution (in DI water) to label the outer shell. Imaging was performed on a Nikon N-STORM super-resolution system equipped with a Hamamatsu ORCA-Quest qCMOS camera in bioluminescence mode (no illumination, 1 s exposure per frame).

Chemical stimulation was delivered by gently adding 500 µL of either acidic (pH 4) or basic (pH 10) solution to the imaging well; control wells received L1 medium alone. Bioluminescent emission was recorded for 60 s (1 frame/s). DAPI (Ex/Em: 358/461 nm) and Cy5 (Ex/Em: 640/670 nm) fluorescence channels were acquired pre- and post-stimulation to confirm cell positioning and viability. To quantify bioluminescent activation patterns over time, single-cell image sequences were analyzed using a custom pipeline in MATLAB and ImageJ. Raw frames were background-subtracted and split into DAPI (cell mask) and luminescence (signal) channels. Thecal shell borders were segmented via thresholding and binarization of DAPI fluorescence to generate cell-specific masks. For each cell, bioluminescent frames were aligned by timepoint and summed temporally to generate cumulative emission maps. These integrated bioluminescence projections were overlaid with corresponding cell masks to compute signal coverage as a percentage of the total cell area:

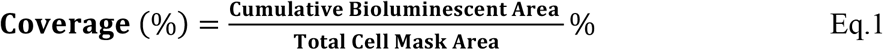

Analysis was conducted across *N* = 10 cells per condition.

### Hydrogel Fabrication and Morphological Characterization

Hydrogel beads were fabricated by dispensing 5 µL droplets of 4 wt% sodium alginate into a 1 wt% CaCl_2_ crosslinking bath and allowing ionic gelation to proceed for 20 min. For cell-laden constructs, *P. lunula* cultures harvested during their dark phase were centrifuged at 1,500 rpm for 10 min, resuspended in sterile L1 medium, and gently mixed into the alginate precursor prior to dispensing, following the same protocol used for acellular beads. For morphological analysis, hydrogel beads were snap-frozen in dry ice, lyophilized overnight, and sectioned. Samples were sputter-coated with a 4 nm titanium layer and imaged under high vacuum using scanning electron microscopy (SEM; Hitachi SU3500) to assess internal microarchitecture.

### Cell Viability within Living Light Materials

Hydrogel constructs encapsulating *P. lunula* at densities of 150, 300, 600 × 10^3^ cells mL^−1^ were formed as described and cultured in L1 medium under standard conditions. Cell viability was assessed using MTS assay (CellTiter 96®, Promega). Individual beads were incubated with MTS reagent (20 µL per well) for 2 h at 24 °C, and absorbance at 490 nm was quantified (Tecan Spark reader).

### 3D Fluorescence Imaging of Living Light Materials

*P. lunula* cells were fixed in 2.5% glutaraldehyde (in PBS, 60 min at room temperature), washed 3 times in PBS, and stained with DAPI (1:1000 dilution in DI water, 15 min). Cells were then encapsulated in 4 wt% sodium alginate at a density of 150 × 10^3^ cells mL^−1^ and crosslinked by dropwise addition in 1 wt% CaCl_2_ solution (20 min at room temperature) to form hydrogel beads. Cross-sectional imaging was performed using an EVOS F500 widefield fluorescence microscope equipped with DAPI (Ex/Em: 357/447 nm) and Cy5 (Ex/Em: 635/692 nm) filter sets using a 4x air objective. For volumetric reconstruction, intact hydrogel beads were imaged using a Bruker TruLive3D light-sheet microscope equipped with dual-channel laser excitation (405 nm for DAPI, 640 nm for Cy5). Image acquisition was conducted using a 25x water-immersion detection objective (NA = 1.1) with a z-step size of 1.0 μm and processed with Bruker LuxBundle software.

### Extrusion-Based 3D Bioprinting of Living Light Materials

Sodium alginate bioinks (4 and 6 wt%) were pre-crosslinked by mixing with 1 wt% CaCl_2_ at defined volumetric ratios (1:0 to 1:1, alginate:CaCl_2_) using a dual-syringe connector to ensure uniform partial gelation. *P. lunula* cells were gently incorporated into the pre-crosslinked alginate (600 × 10^3^ cells mL^−1^ final concentration) immediately prior to printing. Cell-laden inks were loaded into sterile 3 mL cartridges and extruded using a BIO X extrusion-based bioprinter (Cellink) fitted with a 22G conical nozzle (inner diameter: 410 µm) at ambient temperature. Constructs were printed onto glass slides or plastic supports and subsequently post-crosslinked by immersion in 1 wt% CaCl_2_ for 10 min. After crosslinking, all constructs were rinsed with sterile L1 medium and incubated in fresh L1 at 24 °C under standard circadian conditions (12 h light/12 h dark) until further analysis.

### Rheological Characterization of Bioinks

Hydrogel rheology was assessed using an Anton Paar MCR 702e rheometer with 25 mm parallel plates (gap: 0.5 mm). Shear rate sweep (0.01–10% strain at 1 Hz) and viscosity measurements (0.001 to 100 s^−1^) were performed at 24 °C to determine viscoelastic moduli (G′, G″) and shear-thinning behavior.

### Chemical Stimulation and Imaging of Living Light Materials

To evaluate single-cell bioluminescent responses within living hydrogels, cell-laden constructs (4 wt% alginate, 300 × 10^3^ cells mL^−1^) were cast as 10 µL thin films, ionically crosslinked in 1 wt% CaCl_2_ for 20 min, and incubated in L1 medium. Prior to imaging, cells were stained with DAPI (1:1000 dilution, 50 µL), followed by the addition of 500 µL of stimulation buffer (pH 4, pH 10, or L1 medium as control) applied directly to the hydrogel surface.

Macroscopic bioluminescence of larger constructs was imaged using a Canon EOS 90D DSLR camera (f/1.4, ISO 12,800, 30 fps) following the same stimulation protocol. Videos were processed with custom scripts in MATLAB and ImageJ to extract temporal overlays. Quantitative bioluminescence was assessed using an IVIS Lumina system over a 10-minute window post-stimulation, with total photon counts at 5 minutes recorded for analysis (N = 8).

### Chemo-Mechanical Stimulation and Real-Time Monitoring of Living Light Materials

Cylindrical living light materials (d=10 mm, h=3 mm; N = 8) were preconditioned in pH 4, pH 10, or L1 medium for 2 min, then compressed to 80% strain at 0.2 mm s^−1^ using a universal testing machine (Insight II, MTS Systems) under dark conditions. Bioluminescence was recorded in real-time using a Canon EOS 90D camera. Synchronization and intensity quantification were performed with custom MATLAB and Python scripts, extracting mean grayscale values from ROI across frames and aligning with force-displacement data.

### Longitudinal Bioluminescence and Viability Assessment

Hydrogel beads (300 × 10^3^ cells mL^−1^) were subjected to weekly stimulation over a four-week period using either pH 4 buffer, pH 10 buffer, or L1 medium as a control (500 µL per well; N = 8). Bioluminescence was monitored using IVIS imaging for 10 minutes following each stimulation. Hydrogel beads were then transferred to fresh medium and cultured under standard conditions until the subsequent stimulation cycle.

The fraction of luminescent-positive constructs was used to generate Kaplan–Meier-like survival curves representing functional longevity. Cell viability was qualitatively assessed via chlorophyll autofluorescence (Cy5 channel) using an EVOS F5000 fluorescence microscope.

## Statistical Analysis

All data are presented as mean ± standard deviation unless otherwise stated. Group comparisons were performed using two-way ANOVA followed by Tukey’s post hoc test. One-way ANOVA was used for single-factor analyses. Longitudinal functional data were assessed using Kaplan–Meier survival analysis. Statistical significance was defined as *P* < 0.05. All statistical analyses and visualizations were conducted using GraphPad Prism (v9.0) and MATLAB (R2023a).

## Authors Contributions

The grant funding this work was awarded to W.S. G.B. and W.S. conceived, designed, and supervised the study. G.B. established the methods for chemical stimulation for triggering bioluminescence in living materials. G.B. developed experimental protocols and performed experiments for chemical stimulation, hydrogel formulation and 3D printing, fluorescence microscopy, cellular viability assays, and IVIS imaging. J.M. conducted N-STORM Imaging. Light sheet microscopy was performed by J.M. C.P.L., J.M. and G.B. optimized formulation printability. C.P.L. and G.B. carried out 3D printing and chemical-mechanical stimulation experiments. G.B. and C.P.L. performed rheological characterization. J.E.E. and G.B. performed SEM imaging. Data analysis was conducted by G.B. and C.P.L. The manuscript was written by G.B. with input from all co-authors.

## Funding

This research was made possible by the Department of Civil, Environmental, and Architectural Engineering, the Materials Science and Engineering Program, the College of Engineering and Applied Sciences, and the Living Materials Laboratory at the University of Colorado Boulder. This project is supported by Schmidt Sciences, LLC. This research was also supported in part by the Colorado Shared Instrumentation in Nanofabrication and Characterization (COSINC-CHR (Characterization), College of Engineering & Applied Science, University of Colorado Boulder) and the Biofrontiers Advanced Light Microscopy Core (ALM Core, Biofrontiers Institute, University of Colorado Boulder). The authors would like to acknowledge the support of Mr. Matthew H. Fyfe for logistical and technical assistance throughout this research. We are grateful to Drs. Joe Dragavon and Evolène Premilieu of the BioFrontiers Advanced Light Microscopy Core for training and support with light-sheet and N-STORM imaging. We also thank Dr. Adrian Gestos (CU Boulder COSINC) for SEM training, and Aseem Visal and Prof. Carson Bruns for training and access to rheological instrumentation.

## Notes

### Competing Interest Statement

The authors have declared no competing interest.

