## Supplementary Figures for "Chemical Stimulation Sustains Bioluminescence of Living Light Materials"

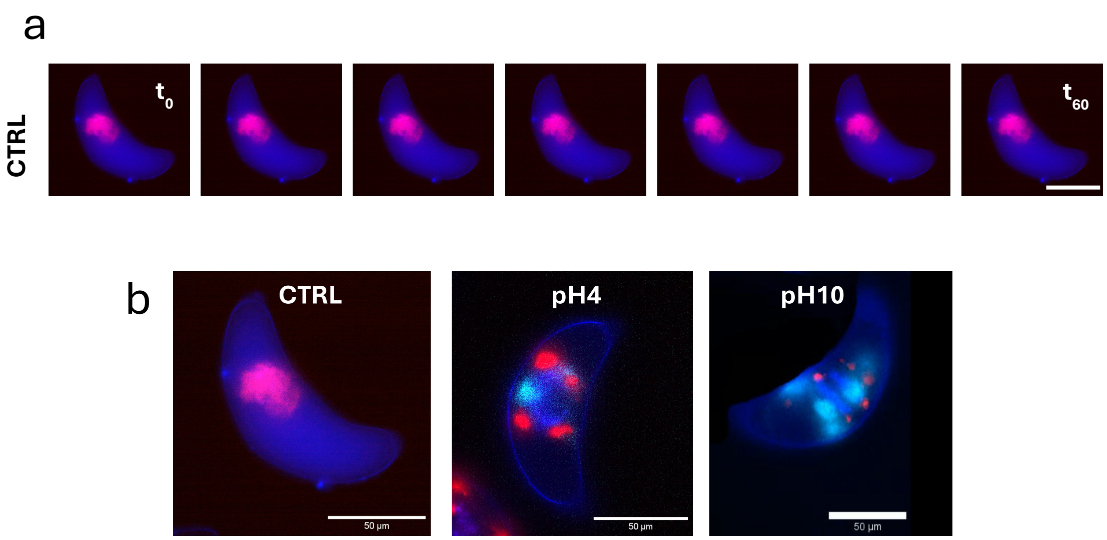


**Figure S1 | Absence of bioluminescent activity in untreated Pyrocystis lunula cells.**

a, N-STORM time-lapse imaging of control (untreated) P. lunula cells reveals no detectable bioluminescent signal over the observation period, confirming the necessity of chemical stimulation to elicit light emission. Scale bars, 50 µm.

b, Representative N-STORM videos (2.5× speed) comparing bioluminescent responses under control, acidic (pH 4), and basic (pH 10) conditions. Scale bars, 50 µm.

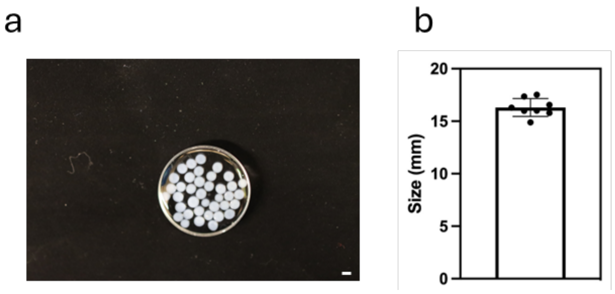


**Fig. S2 | Size distribution of alginate beads following ionic crosslinking.**

a, Brightfield image of ionically crosslinked alginate beads (scale bar = 1 mm). b, Quantification of bead diameter (1.6 ± 0.1 mm , n = 8) confirms consistent size formation across the batch.

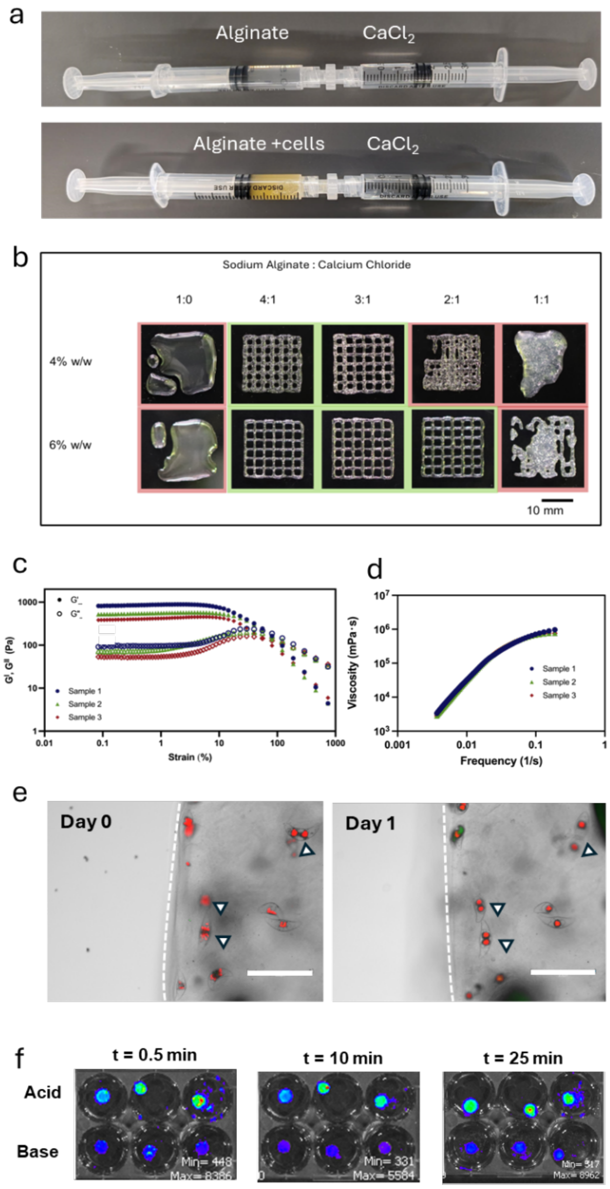


**Figure S3 | Pre-crosslinking strategy for enhanced printability of *P. lunula*–laden alginate hydrogels.**

**a**, Coaxial syringe-based mixing of sodium alginate and $CaCl_2$ enables partial ionic pre-crosslinking, improving rheological properties for controlled extrusion. Top: alginate-only formulation; bottom: alginate with embedded *P. lunula* cells.

**b**, Brightfield images of bioink formulations evaluated for shape fidelity post-printing. Printability indicated by green borders; red outlines mark suboptimal conditions. Scale bar, 10 mm.

**c**, Rheological analysis of optimized formulations (4 wt% alginate, 4:1 alginate:$CaCl_2$) highlighting storage modulus and **d,** shear-thinning behavior, consistent with extrusion-based bioprinting (n = 3).

**e**, Brightfield and Cy5 fluorescence overlays showing spatial confinement of *P. lunula* and post-printing viability at day 0 and day 1. Arrows indicate proliferation. Scale bars, 300 µm

**f**, IVIS imaging of P. lunula–laden hydrogels shows sustained bioluminescence up to 25 minutes post-stimulation under acidic conditions (n = 3).

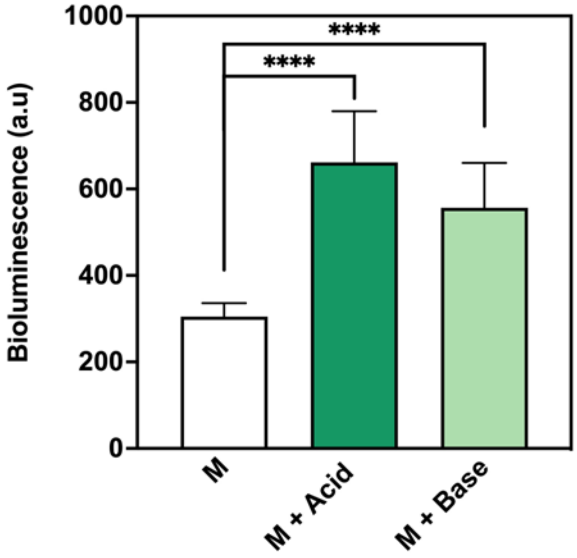


**Figure S4 | Synergistic enhancement of overall bioluminescence by combined chemical and mechanical stimulation.**

Quantification of total emitted bioluminescence (a.u.) from *P. lunula*–laden constructs subjected to mechanical stimulation (M) alone or following acid or base preconditioning (M + Acid, M + Base). (n = 8, mean ± s.d.; ****$P < 0.0001$, two-way ANOVA with Tukey's post hoc test).
